# Transcriptome assembly from long-read RNA-seq alignments with StringTie2

**DOI:** 10.1101/694554

**Authors:** Sam Kovaka, Aleksey V. Zimin, Geo M. Pertea, Roham Razaghi, Steven L. Salzberg, Mihaela Pertea

## Abstract

RNA sequencing using the latest single-molecule sequencing instruments produces reads that are thousands of nucleotides long. The ability to assemble these long reads can greatly improve the sensitivity of long-read analyses. Here we present StringTie2, a reference-guided transcriptome assembler that works with both short and long reads. StringTie2 includes new computational methods to handle the high error rate of long-read sequencing technology, which previous assemblers could not tolerate. It also offers the ability to work with full-length super-reads assembled from short reads, which further improves the quality of assemblies. On 33 short-read datasets from humans and two plant species, StringTie2 is 47.3% more precise and 3.9% more sensitive than Scallop. On multiple long read datasets, StringTie2 on average correctly assembles 8.3 and 2.6 times as many transcripts as FLAIR and Traphlor, respectively, with substantially higher precision. StringTie2 is also faster and has a smaller memory footprint than all comparable tools.

## Introduction

Measuring the abundances of transcripts in an RNA-sequencing (RNA-seq) dataset is a powerful way to understand the workings of a cell. Simply aligning reads to a reference genome can provide rough estimates of the average expression of genes and hint at differential use of splice sites (Li and Dewey 2011), but to create an accurate picture of gene activity, one must assemble collections of reads into transcripts. Alternative splicing is very common in eukaryotes, with an estimated 90% of human multi-exon protein-coding genes and 30% of non-coding RNA (ncRNA) genes having multiple isoforms (Wang, Sandberg et al. 2008, Cabili, Trapnell et al. 2011). While the number of annotated human protein coding genes has remained more or less constant over the last decade, the number of ncRNA genes and protein coding isoforms has continued to increase (Pertea, Shumate et al. 2018).

Second-generation sequencers, such as those from Illumina, can produce hundreds of millions of short (∼100bp) RNA-seq reads. Reads of this length usually span no more than one intron, except in rare cases of very small exons. By assembling the short reads, we can reconstruct full-length transcripts and identify novel genes and gene isoforms. There are two main approaches to transcriptome assembly: *de novo* and reference-guided. *De novo* transcriptome assemblers such as Trinity (Grabherr, Haas et al. 2011) and Oases (Schulz, Zerbino et al. 2012) find overlaps between reads and attempt to chain them together into full transcripts. This is complicated by the presence of paralogous genes and transcripts with many isoforms that largely overlap one another, and as a result this approach produces highly fragmented and error-prone transcriptomes. Reference-guided assemblers such as Cufflinks (Trapnell, Williams et al. 2010), Bayesembler (Maretty, Sibbesen et al. 2014), StringTie (Pertea, Pertea et al. 2015), TransComb (Liu, Yu et al. 2016), and Scallop (Shao and Kingsford 2017) take advantage of an existing genome to which the RNA-seq reads are first aligned using a spliced aligner such as HISAT (Kim, Langmead et al. 2015) or STAR (Dobin, Davis et al. 2013). These assemblers can then build splice graphs (or other data structures) based on the alignments, and then use those graphs to construct individual transcripts. Some reference-guided assemblers can also use the exon-intron annotation of known transcripts as an optional guide, allowing them to favor known genes where possible. A recent study (Voshall and Moriyama 2018) found that StringTie outperforms both Cufflinks and Bayesembler, by assembling more correctly transcripts and at a higher precision, while the original Scallop study (Shao and Kingsford 2017) showed that on some datasets, Scallop can achieve higher sensitivity and precision than StringTie (version 1.3) and TransComb.

StringTie and other transcriptome assemblers estimate transcript abundance based on the number of aligned reads assigned to each transcript. More recently, alternative methods such as Sailfish (Patro et al. 2014), Salmon (Patro, Duggal et al. 2017) and Kallisto (Bray, Pimentel et al. 2016) demonstrated that one can estimate abundances by assigning reads to known transcripts based on exact k-mer matching, which produces dramatic gains in speed by dropping the requirement for precise base-level read alignment. However, these alignment-free methods are not able to detect novel genes or isoforms, and they show poorer performance in quantifying low-abundance and small RNAs compared to alignment-based pipelines (Wu, Yao et al. 2018).

The first release of StringTie proposed a method to use a limited version of *de novo* transcriptome assembly via the construction of super-reads, which were originally developed for whole-genome assembly (Zimin, Marcais et al. 2013). Conceptually, super-reads are constructed by extending each end of a short read as long as there is a unique extension based on a k-mer lookup table. This creates a collection of synthetic long reads with the low error rate of short reads. Because they are longer, they are more likely to align uniquely to the genome, which in turn might simplify the splice graph of a gene. Super-reads were used in a limited capacity in StringTie 1.0, only filling in the gap between paired-end reads. In that limited implementation, a super-read was used to replace a pair of reads, allowing it to be treated like a single, unpaired read. One difficulty in using super-reads is that the algorithm used to create them for genome assembly includes an error correction step, which in the context of RNA-seq assembly could over-write k-mers from low-abundance transcripts. Another complication is that a full super-read may contain many short reads, and thus it cannot be counted as a single read during the quantification step. We have therefore developed an expectation-maximization (EM) algorithm to distribute read coverage between super-reads.

While second-generation sequencers produce very large numbers of reads, their read lengths are typically quite short, in the range of 75-125 bp for most RNA-seq experiments. These short reads often align to more than one location, and we designate such reads as “multi-mapping”. Short reads also suffer the limitation that they rarely span more than two exons, making the splice graph difficult and sometimes impossible to traverse accurately for genes with multiple exons and many diverse isoforms, no matter how deeply they are sequenced. These issues can be alleviated by third-generation sequencing technologies such as those from Pacific Biosciences (PacBio) and Oxford Nanopore Technologies (ONT). These long-read technologies, which can produce read lengths in excess of 10,000 bp, have dramatically improved whole-genome assemblies (Zimin, Puiu et al. 2017), and when used for RNA-seq experiments, they offer the potential for large gains in the accuracy of isoform identification and discovery (Au, Sebastiano et al. 2013, Tilgner, Grubert et al. 2014, Kuosmanen, Norri et al. 2018). While some reads produced by third-generation sequencers cover the full length of RNA transcripts, many will inevitably capture only partial transcripts. This happens for a variety of reasons; e.g., (1) RNA degrades quickly and may be shorter than full length by the time it is captured for sequencing; (2) long molecules can break during library preparation; or (3) in cDNA sequencing, the reverse transcription step may fail to capture the full RNA molecule. Thus computational tools that only consider reads which fully cover a transcript will be forced to discard many reads, possibly causing a substantial reduction in sensitivity. To date, though, long reads have not been widely adopted for transcriptome assembly, in part because they have a much higher error rate (typically 8-10% or higher), making alignment difficult (Roberts, Carneiro et al. 2013, Jain, Fiddes et al. 2015), and also because long-read sequencers have much lower throughput, which makes quantification of all but the highest-expressed genes impossible.

Various tools have recently been developed to correct errors and/or extract full-length transcripts from genome alignments of long RNA-seq reads. TranscriptClean (Wyman and Mortazavi 2019) corrects mismatches, indels, and non-canonical splice-sites in long-read alignments, but does not attempt to identify full-length transcripts. FLAIR (Tang et al. 2018) corrects splice-sites based on a known annotation and outputs transcripts from the annotation that are fully covered by “high-confidence” reads. An alternative to these approaches, which rely solely on known transcripts, one can assemble long-read fragments using the same methods used for short-read transcriptome assembly. In addition to finding novel transcripts, the assembly approach can more readily handle fragments that match multiple isoforms, and it can correct alignment errors by forming a consensus from multiple reads. Traphlor (Kuosmanen et al. 2016) is the only previously system designed to assemble high-error long reads, although we show it performs very poorly on both simulated and real data.

Here we present StringTie2, a major new release of the StringTie transcript assembler, which is capable of assembling both short and long reads, as well as full-length super-reads. Our results on 33 Illumina RNA-seq datasets demonstrate that StringTie2 is more accurate than Scallop, the next-best performing transcriptome assembler of those currently available. The use of super-reads also consistently improves both the sensitivity and precision of StringTie2 assemblies. When applied to long reads, StringTie2 assembles the reads substantially more accurately, faster, and using less memory than FLAIR, the next-best performing tool for long-read analysis. As opposed to FLAIR, StringTie2 can also identify novel transcripts from the long-read data, even when no reference annotation is provided.

## Results

### Transcriptome assembly of short RNA-seq reads

We first used simulated data to compare the sensitivity and precision of StringTie2, with and without super-reads, to that of Scallop (Fig. 1). We define sensitivity as the percent of expressed transcripts that match a transcript predicted by each tool, and precision as the percent of predicted transcripts that match an expressed transcript. (Note that precision is equivalently called positive predictive value.) We say two transcripts match if they have identical intron chains and their first and last exons begin and end within 100bp of each other. We tuned the default parameters of StringTie2 to have approximately the same precision as StringTie (version 1.3) on this simulated data. StringTie2, with default parameters, is more sensitive and precise than Scallop on simulated data, and the use of super-reads increases both the sensitivity and precision of StringTie2 compared to using short-read alignments alone. We also computed the Spearman correlation coefficients of the expression levels predicted by each tool compared to the true expression levels on simulated data (Table 1). StringTie2 has a higher correlation than Scallop over all predicted and expressed transcripts, and the use of super-reads improves this correlation further.

**Table 1.** Spearman correlation coefficients for the simulated short-read assemblies. “Spearmen Predicted” only includes transcripts that each tool assembled. For non-assembled transcripts in “Spearman all” the predicted expression was set to zero.

|  | **Spearman**<br>**All** | **Spearman**<br>**Predicted** |
| --- | --- | --- |
| **Scallop** | 0.726 | 0.828 |
| **StringTie2** | 0.781 | 0.925 |
| **StringTie2+SR** | 0.788 | 0.930 |

**Figure 1.**
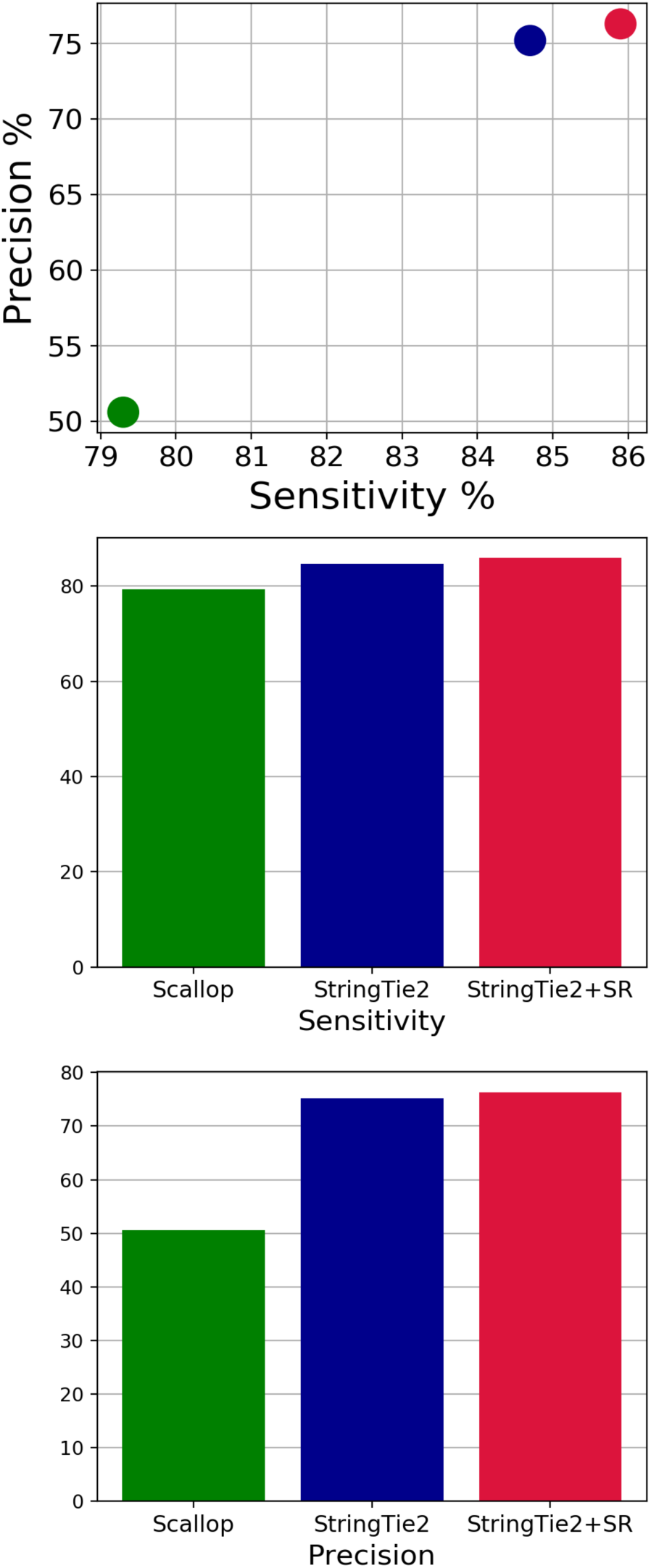
Sensitivity and precision of Scallop, StringTie2, and StringTie2 with super-reads (StringTie2+SR) on simulated human short-read data, containing 150 million 75-bp paired-end reads. Only those transcripts that were completely covered by input reads are considered.

We next evaluated performance on real short-read data, which is considerably more complex than simulations. For this data, we cannot know with certainty which transcripts were expressed in each dataset, nor can we know their precise expression levels. However, it is generally safe to assume that an assembler is more sensitive if it assembles more transcripts matching known annotations (i.e., transcripts from a published database of known genes), and that it is more precise if the known transcripts represent a higher proportion of all the transcripts that are output by the assembler. Therefore we define sensitivity and precision using the union of all annotated transcripts correctly predicted by each tool on a given sample as our set of “expressed” transcripts for that sample (also see Methods).

We ran StringTie2 and Scallop on 23 short-read datasets from human, five from *Arabidopsis thaliana*, and five from *Zea mays* (see Methods). On all datasets, StringTie2 was more accurate than Scallop (Supplementary Table S1). Figure 2 shows that StringTie2 improved on both sensitivity and precision when compared to Scallop, with an average increase of 3.9% and 47.3% in sensitivity and precision respectively. StringTie2 outperformed Scallop on both metrics on all but one plant dataset. Scallop had a slightly higher precision than Stringtie2 (17.4% vs 16.3%, respectively) on one *Z. mays* dataset (ERR986144), however even on that sample, StringTie2 still obtained a 24% relative increase in sensitivity (see Fig. 2).

**Figure 2.**
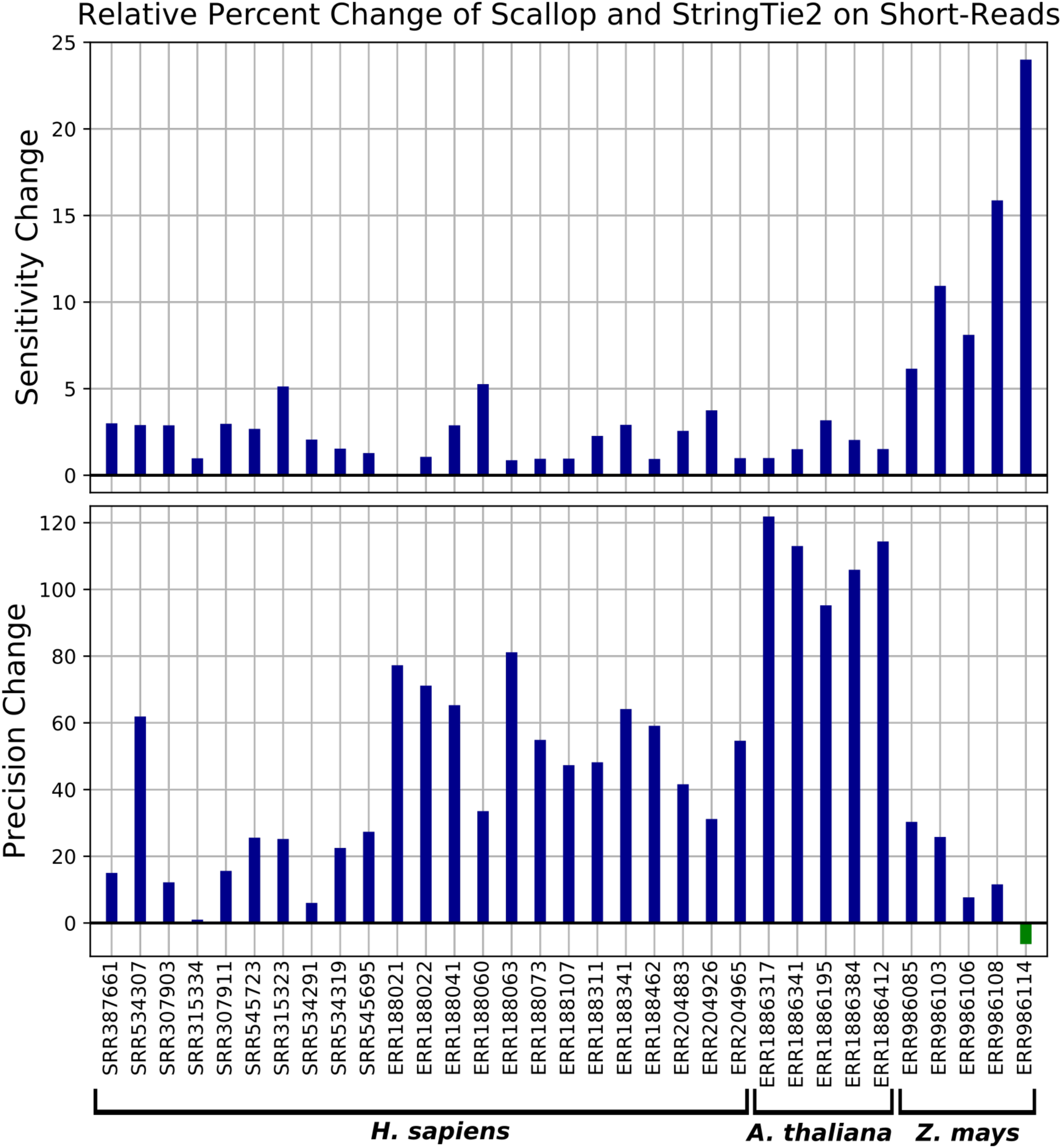
Relative change in sensitivity and precision of StringTie2 vs. Scallop on real short-read data from humans, *Arabidopsis thaliana*, and *Zea mays*.

After inspecting the read alignments of the *Z. mays* datasets in IGV (Robinson et al. 2011), we noticed that many of the expressed transcripts appeared to be fragmented, because the read alignments did not cover the full length of the transcripts; i.e., there were gaps in coverage. Although StringTie2 assembled those fragments correctly, its precision was hurt because its output did not contain full-length transcripts. Both StringTie2 and Scallop have a parameter that allows changing the maximum allowed gap between two read alignments that the tools considers to be part of the same transcript. We increased this parameter from the default 50bp to 200bp for both tools. We also disabled trimming of the transcripts assembled by StringTie2; this is triggered when the read coverage drops below a given threshold at the 5’ or 3’ ends of a transcripts. When this occurs, StringTie2 doesn’t extend the transcripts past the drops in coverage. We could not find a similar parameter in Scallop. With these parameter changes, StringTie2 has both substantially higher sensitivity and precision than Scallop on all *Z. mays* datasets, including ERR986144 (Supplementary Table. S2). Additionally, StringTie2’s sensitivity and precision are substantially increased after this parameter adjustment, compared to the runs with default parameters.

StringTie2 is not only more accurate than Scallop, but also more time and memory efficient. Averaging over all real short-read datasets, StringTie2 runs 1.8 times faster than Scallop and uses 17 times less memory (Supplementary Table S3).

The use of super-reads increases both the sensitivity and precision of StringTie2 on all human datasets and all but three plant datasets (Fig. 3). Of the latter three, StringTie2 had an increase in precision but no change in sensitivity on one *Z. mays* dataset, and an increase in sensitivity but no change in precision on two *Arabidopsis* datasets.

**Figure 3.**
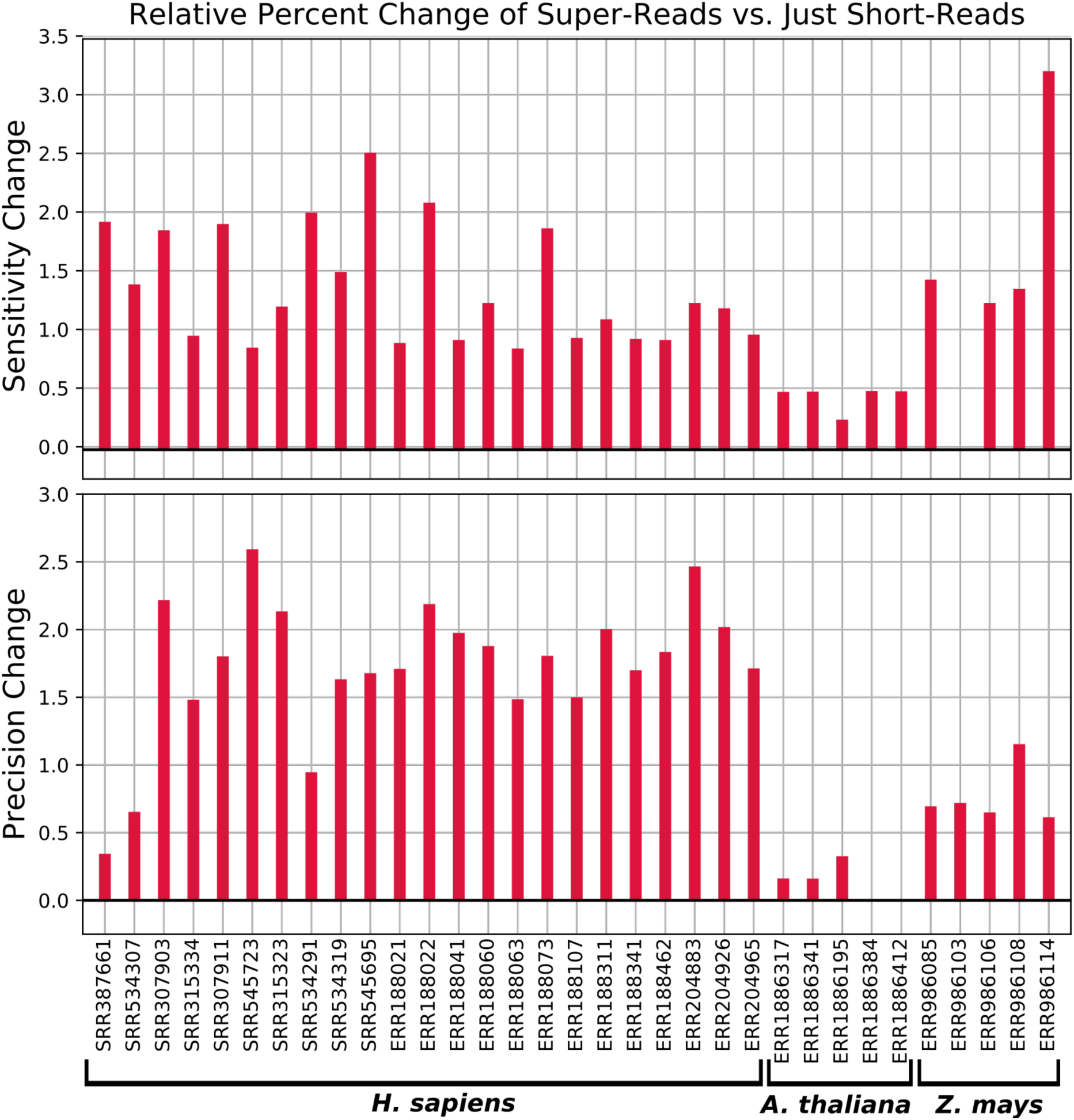
Relative change in percent sensitivity and precision when using super-reads with real short-read data.

### Transcriptome assembly of third-generation RNA-seq long-reads

We next compared StringTie2’s performance on long-reads with that of FLAIR and Traphlor, the only other systems that can process long-read RNA sequencing data. Because we cannot know the true transcript in real RNA-seq data sets, we first used simulated data to assess the accuracy of all tested tools. We obtained five simulated datasets generated by Krizanovic et al. (2017), who used the DNA simulator PBSIM (Ono et al. 2013) tuned to mimic the statistics of either PacBio or ONT RNA-seq data. These datasets consist of a *Saccharomyces cerevisiae* PacBio run, two *Drosophila melanogaster* runs (one PacBio, one ONT), and two human chromosome 19 runs (one PacBio, one ONT). We ran StringTie2, FLAIR, and Traphlor on these simulated datasets and computed sensitivity and precision as before (Fig. 5). FLAIR requires gene annotation as a guide to alignment, so we also ran StringTie2 with the same guide annotation in order to make a direct comparison.

**Figure 4.**
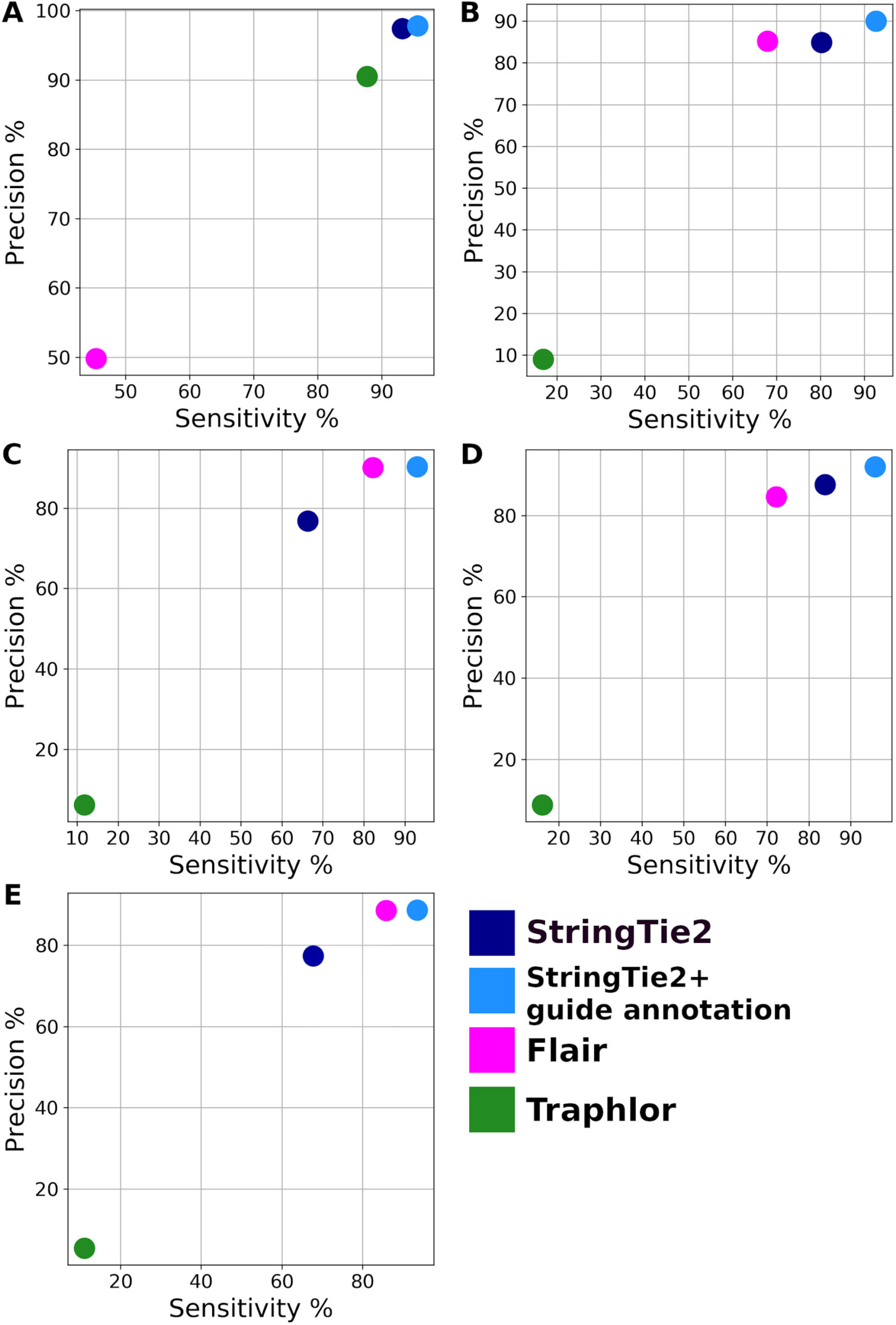
Sensitivity and precision of StringTie2 (with and without guide annotation), FLAIR, and Traphlor on long read simulated data from A) PacBio *Saccharomyces cerevisiae*, B) PacBio *Drosophila melanogaster*, C) PacBio *Homo sapiens*, D) ONT *D. melanogaster*, and E) ONT *H. sapiens.*

**Figure 5.**
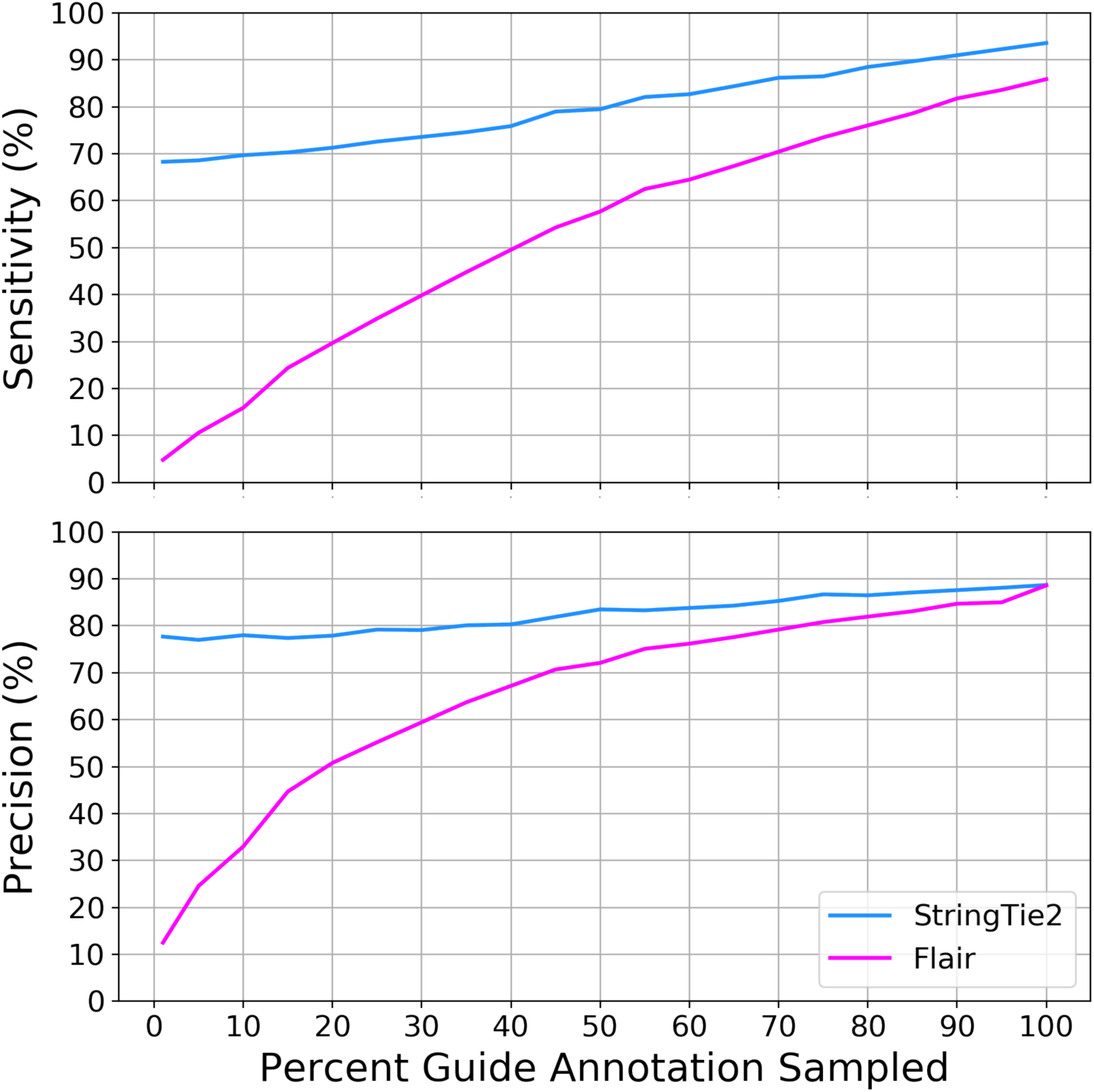
Sensitivity and precision of StringTie2 and FLAIR running on ONT human chromosome 19 simulated data using random samples of different sizes of the human chromosome 19 annotation as a guide.

Traphlor had lower sensitivity and precision than StringTie2 and FLAIR on all datasets except for the *S. cerevisiae* PacBio data, where it outperformed FLAIR on both metrics (Fig 5a). StringTie2 with a guide annotation outperformed FLAIR on all datasets, and in some cases StringTie2 *without* a guide annotation performed equally well. Because this was simulated data, the guide annotation included all transcripts that were present in the sample, even if not all of them were expressed. Real datasets are likely to contain unannotated transcripts and may lack many known, annotated genes entirely.

To demonstrate the performance of each tool when transcripts are missing from the guide annotation, we ran StringTie2 and FLAIR on the human chromosome 19 ONT data using random samples of the chromosome 19 annotation, which we varied to contain from 1% to 100% of the transcripts. Results are shown in Figure 5. The sensitivity and precision of FLAIR decreases rapidly as the amount of annotation is reduced; e.g., when only 20% of the annotation is provided, FLAIR’s sensitivity and precision dropped to 30% and 50% respectively. In contrast, with that same amount of annotation, Stringtie2’s results were far better, 74% and 80%. This result demonstrates FLAIR’s strong reliance on the guide annotation and StringTie2’s contrasting ability to assembly transcripts that are not present in the annotation.

We next ran StringTie2, FLAIR, and Traphlor on eight real human long-read datasets: three PacBio datasets enriched for full-length transcripts (PacBioFL), three PacBio datasets containing transcript fragments (PacBioNFL), one nanopore cDNA dataset (NPcDNA), and one nanopore direct RNA-seq dataset (NPDirect). Traphlor failed to produce any transcripts on the NPcDNA dataset, and had drastically worse precision and sensitivity compared to StringTie2 on all other datasets (Supplementary Table S4). Averaging across all datasets on which Traphlor was able to run, StringTie2 correctly assembled 9564 transcripts, 2.6 times more than Traphlor’s 3708 correct assemblies. Compared to FLAIR, StringTie2 with guide annotation correctly identified 16,000 more transcripts on average, with precision that ranged from about three to six times higher (Fig. 6, Supplementary Table S4). FLAIR performed the best on the nanopore direct RNA-seq dataset where it correctly identified 4,442 from the annotation. By comparison, StringTie2 correctly assembled 29,744 transcripts, 6.7 times more than FLAIR. Even without using guide annotation, StringTie2 substantially outperformed FLAIR on all of the real datasets (Supplementary Table S4). StringTie2 with annotation is 68 times faster than FLAIR and uses 9 times less memory, averaged over all real long-read datasets. Without annotation, StringTie2 is 93 times faster than FLAIR and uses 27 times less memory (Supplementary Table S5).

**Figure 6.**
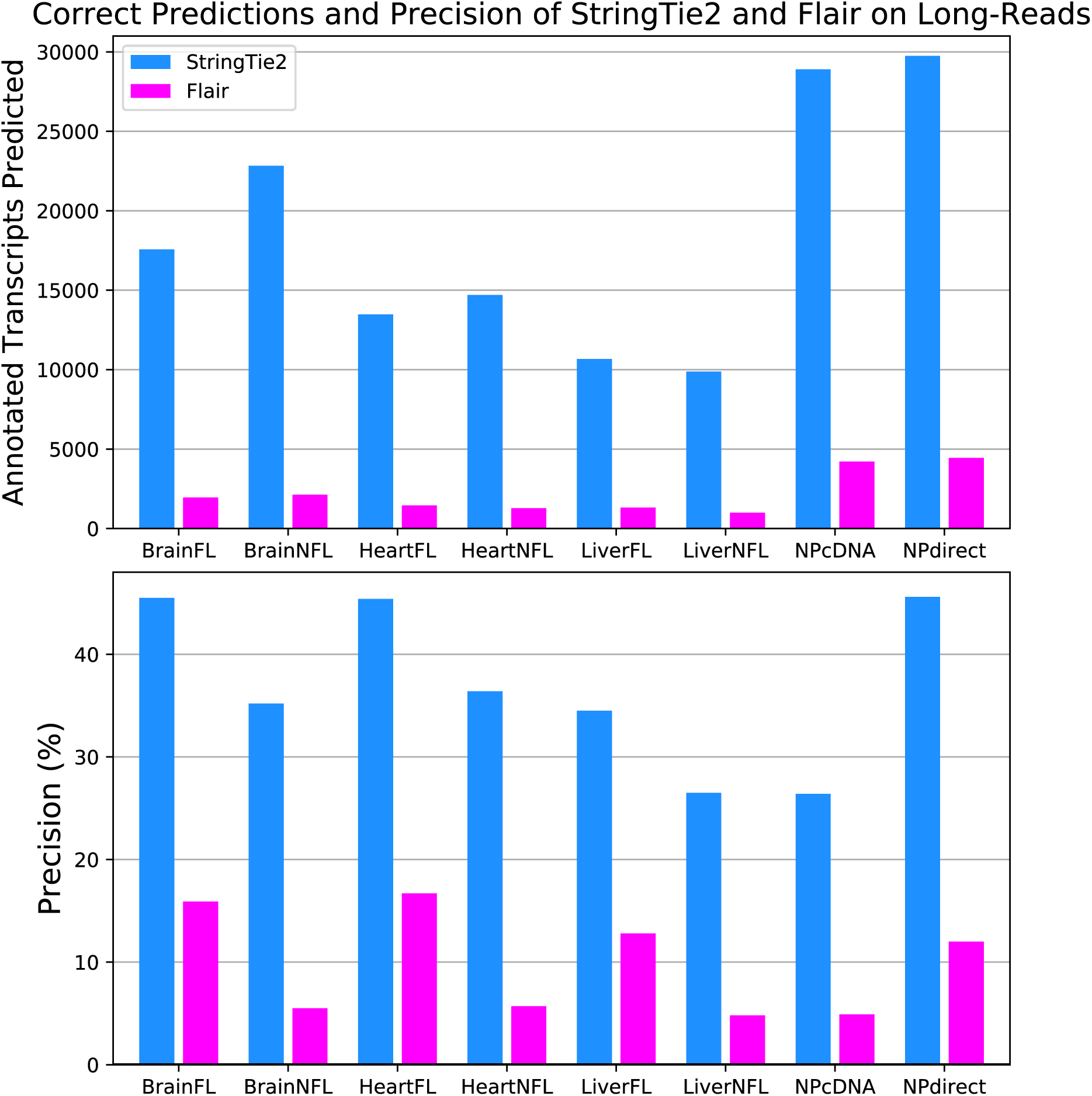
Number of correctly assembled transcripts correctly and precision of StringTie2 and FLAIR on real PacBio (FL = full length, NFL = not full length) and ONT (NP direct RNA and cDNA) human data. For both plots, any transcripts assembled by either tool were considered correct if they exactly matched all introns from a known, annotated transcript.

## Discussion

Our results show that StringTie2 is more accurate than Scallop on both real and simulated data sets. On all short-read data sets evaluated here, StringTie2 obtained better sensitivity, higher quantification accuracy, faster runtime, and lower memory usage than Scallop. Close inspection of the only sample where StringTie2 did not have higher precision (ERR986114, Fig. 2) showed that there were many gaps between the read alignments within individual transcripts, possibly due to the repetitive nature of the *Zea mays* genome. Scallop obtains a slightly higher precision on that data, but assembles many fewer transcripts than StringTie2. After making a small parameter adjustment as described above, StringTie2 outperformed Scallop on all metrics on all *Zea mays* data sets (Supplementary Table S1). This demonstrates how default parameters may not be optimal for all datasets, and that more careful parameter selection can improve assembly regardless of the tool used. The same point was recently demonstrated specifically for Scallop and StringTie by DeBlasio and Kingsford (2018).

The use of super-reads introduces partial *de novo* assembly into StringTie2, which we have shown improves sensitivity, precision, and abundance estimation on real and simulated data.

The high error rates of long reads generated by third-generation sequencers present distinct challenges that make identifying the exact exon-structure of a transcript difficult. Alignments of high-error long reads generated from the same locus usually disagree with one another, particularly surrounding splice sites (Supplementary Figures S1, and S2). They also often disagree about the presence or absence of particular exons, especially if the exons are small (Supplementary Figures S2, and S3). The results shown here demonstrate that StringTie2 is the most accurate method for assembly of transcripts from long, high-error rate reads. This has the potential to greatly improve the sensitivity of analyses using long-read RNA-seq data, which in the past has relied primarily on reads that span transcripts end-to-end. The built-in consensus calling in StringTie2 should also lessen the need for a separate error correction step from tools such as TranscriptClean (Wyman and Mortazavi 2019). In addition to its fast runtime and small memory footprint, StringTie2 requires no dependencies and can be easily run as a single command, unlike tools such as FLAIR which consist of a series of scripts that can each fail independently. It is also multi-threaded, which allows it to be run in parallel on multi-processor computers and can significantly reduce the “wall clock” time of assembly.

Further development of long-read RNA-seq technologies will increase the usefulness of StringTie2. In the case of ONT reads, improvements to basecalling will improve alignment quality, which will further improve StringTie2’s assembly. As third-generation sequencers increase their throughput, researchers will also be able to use long-read RNA-seq for accurate transcript-level quantification, which currently requires the higher throughput of short read (i.e., Illumina) sequencers. ONT direct RNA sequencing has additional unique capabilities which are only beginning to be explored, such as the ability to identify RNA base modifications and secondary structure from the raw signal (Workman, Tang et al. 2018). Better transcriptome assemblies will aid these efforts because these read-level features can then be associated with the full transcripts.

## Methods

### RNA-seq real data sets

Ten of the short-read RNA-seq datasets used here were also used by Shao and Kingsford in their evaluation of Scallop (Shao and Kingsford 2017): SRR307903, SRR315323, SRR315334, SRR534307, SRR545723, SRR307911, SRR387661, SRR534291, SRR534319, and SRR545695. Three of these ten–SRR534291, SRR534319, and SRR545695–were used in the original StringTie study (Pertea, Pertea et al. 2015), which described StringTie1. We also examined 13 short-read samples randomly selected from the GEUVADIS dataset (Lappalainen, Sammeth et al. 2013): ERR188021, ERR188022, ERR188041, ERR188060, ERR188063, ERR188073, ERR188107, ERR188311, ERR188341, ERR188462, ERR204883, ERR204926, and ERR204965. Five *Arabidopsis* datasets were obtained from (James, Syed et al. 2012): ERR1886195, ERR1886317, ERR1886341, ERR1886384, and ERR1886412. Five *Z. mays* datasets were obtained from (Wang, Tseng et al. 2016): ERR986085, ERR986103, ERR986106, ERR986108, and ERR986114. The “full length” and “not full length” PacBio datasets were downloaded from http://datasets.pacb.com.s3.amazonaws.com/2014/Iso-seq_Human_Tissues/list.html. The ONT direct RNA-seq and cDNA datasets are from the NA12878 RNA sequencing consortium (Workman, Tang et al. 2018).

### Reference Genomes and Annotations

All human RNA-seq reads were mapped to the main chromosomes of hg38, i.e. not including the “alternate” and “random” scaffolds. The annotation used to compute the accuracy of transcriptome assemblies and to create the human short-read simulated data and the annotation guided assemblies contains all full-length protein and long non-coding RNA transcripts from RefSeq, release GRCh38.p8. The *A. thaliana* RNA-seq reads were aligned to the TAIR10 assembly, and the full corresponding annotation was used for determining accuracy (Lamesch et al. 2011). The *Z. mays* reads were aligned to the B73 RefGen assembly, and the full corresponding annotation was obtained from MaizeGDB (Portwood et al. 2018).

### Simulated Data

A short-read RNA-seq dataset containing 150 million 75-bp paired-end reads was generated using Flux Simulator (Griebel, Zacher et al. 2012) with all protein coding and lncRNA transcripts on the main chromosomes of hg38. The parameters for the simulation were the ones recommended for *H.sapiens* in Supplementary Table S3 from Griebel, Zacher et al. 2012. Long read simulated data for Saccharomyces cerevisiae S288 (baker’s yeast), Drosophila melanogaster r6 (fruit fly), and Homo Sapiens GRCh38.p7 (human) was obtained from Krizanovic et al. (2017). The long reads were simulated using either PacBio (one dataset for yeast, fruit fly, and human each) or MinION ONT profiles (one data set for fruit fly and one for human).

### Alignment and assembly parameters

All short read datasets were aligned using HISAT2 with default parameters. The PacBio and ONT datasets were aligned with minimap2 using the “-splice” option, which enables spliced alignment of noisy long reads. Super-reads were aligned using GMAP because their error profile more closely resembles that of EST sequences, which aligners like minimap2 are not designed for.

The *Z. mays* samples in Supplementary Table S2 were run with the “-t -g 200” options in StringTie2 and the “--min_bundle_gap 200” option in Scallop. StringTie2 was run using the “-L” parameter for all long-read datasets. Three FLAIR sub-commands were run in sequence to obtain the GTF of covered transcripts: “align”, “correct”, “collapse”, using the human reference genome and annotation described above where required. All other assemblies were run using default parameters.

### Accuracy metrics

Similar to Voshall and Moriyama 2018, we used the following metrics to report the accuracy of the transcriptome assemblies:

Sensitivity = TP/(TP+FN)

Precision = TP/(TP+FP)

where TP (or true positives) are correctly assembled transcripts, FP (or false positives) are transcripts that are assembled with errors, and FN (or false negatives) are transcripts in the reference annotation that are missing from the assembly. Sensitivity and precision were determined by running `gffcompare` (Pertea et al. 2016).

Relative percent change in sensitivity (*S*_*r*_) and precision (*P*_*r*_) of StringTie2 versus another method was computed as 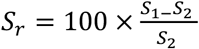, and 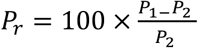, where *S*_1_ and *P*_*1*_ are the sensitivity and precision of StringTie2, and *S*_*2*_ and *P*_*2*_ and the sensitivity and precision of the method which we compare it to (e.g. Scallop or FLAIR). For example, a 10% absolute increase in sensitivity from *S*_2_ = 20% to *S*_1_ = 30% would be reported as a relative increase of 50%.

For real datasets, we have no way to determine exactly what transcripts were truly present in the sample. Therefore, for the purpose of comparison, we defined the set of reference or “true” transcripts to be the union of all annotated transcripts correctly predicted by each tool on a given sample. This metric will overestimate the absolute sensitivity if there are transcripts that no tool predicts, but the relative sensitivity comparison will be accurate because the denominator is the same between samples and therefore cancels out.

### New data structures in StringTie2 compared to StringTie1

StringTie2 builds on our previously developed StringTie1 system, which introduced several key innovations, notably (1) a novel network flow algorithm to reconstruct transcripts and quantitate them simultaneously; and (2) an assembly method to merge read pairs into full fragments in the initial phase (Pertea, Pertea et al. 2015). StringTie2 maintains the same general framework for the assembly and quantification of transcripts but implements much more efficient data structures that overall lead to faster run times and much lower memory usage (see Results). It includes additional techniques designed to handle very long reads, including high error-rate reads produced by the third-generation sequencers, as well as the longer reads that result from the pre-assembly of short reads.

There are three main differences in the way StringTie2 stores aligned reads compared to StringTie1. The first difference is that instead of storing every read individually, StringTie2 collapses reads aligned to the identical location on the genome and keeps a count of how many alignments were collapsed. This simple change has a big impact on the memory required to store input data, because very highly expressed transcripts can sometimes reach a coverage of hundreds of thousands of reads per base. (See for instance Supplementary Figure S4, which illustrates the very high level of expression for the COL1A1 gene in sample SRR534291 that was collected from fetal lung fibroblasts.) However, implementing this change was quite complex, because it also required us to create a different method for storing the pairings between reads, as reads aligned at the same place do not necessarily have their “mates” (the second read in each pair) sharing the same alignments. Previously, for each read StringTie1 stored a pointer to its pair. StringTie2 must instead store an array of pointers to all paired read alignments that are present in the data.

StringTie2 also differs from StringTie1 in its more aggressive strategy for identifying and removing spurious spliced alignments. If a spliced read is aligned with more than 1% mismatches, keeping in mind that Illumina sequencers have an error rate <0.5%, then StringTie2 requires 25% more reads than usual (the default is 1 read per bp) to support that particular spliced alignment. In addition, if a spliced read spans a very long intron (more than 100,000bp), StringTie2 accepts that alignment (and the intron) only if a larger anchor of 25 bp (10bp is the default) is present on both sides of the splice site. Here the term “anchor” refers to the portion of the read aligned within the exon beginning at the exon-intron boundary.

Another improvement in StringTie2 is in its internal representation of its splice graph and of the reads aligned to that graph. Both the assembly of reads into transcripts, as well as the quantification of the resulting transcripts require determining the compatibility between the reads (or fragments) and a path in the splice graph (Pertea, Pertea et al. 2015), which requires many searches of the overlaps between reads and the splice graph. In order to maximize the efficiency of such searches, StringTie1 uses a bit-vector representation of the splice graph, where bits *0* to *n-1* correspond to all nodes in the splice, and bit *n*\**i*+*j* corresponds to a possible edge between nodes *i* and *j* in the splice graph, where *n* is the number of nodes in the graph and *i*<*j* (Supplementary Figure S5). A read or a paired read will therefore be represented by a vector of bits where only the bits that represent the nodes or edges spanned by the read and its pair are set to 1. Because in general many of the nodes in the splice graph are not connected by edges, most bits in this bit-vector representation will be 0; therefore StringTie2 replaces it with a sparse bit-vector data structure, where the bits can only correspond to a node or an edge appearing in the splice graph. Building more efficient data structures in StringTie2 greatly reduced the memory footprint of the StringTie system. On the three datasets from this study that were also examined in the original StringTie release, memory usage was reduced on average by a factor of 40 (Supplementary Figure S6).

### Assembly of long RNA-seq reads

Third-generation sequencing technologies (i.e., from PacBio and Oxford Nanopore sequencing instruments) have an error profile that consists mostly of insertion and deletions, as opposed to second-generation errors that are mostly substitutions. Insertion and deletions are harder to correct than substitutions, and the accuracy of methods for correcting them is generally low (Allam, Kalnis et al. 2015). Further complicating matters, aligning long reads correctly around splice sites is challenging, and mis-alignments lead to spurious edges in the splice graph, which in turn leads to incorrect transcript predictions (Kuosmanen, Norri et al. 2018).

To handle the high error rates in the long reads, we implemented two new techniques in StringTie2. First, we correct potentially wrong splice sites by checking all the splice sites present in the alignment of a read with a high-error alignment rate. If a splice site is not supported by any low-error alignment reads then we try to find a nearby splice site (within 10bp, by default) that is supported by the most alignments among all nearby splice sites. If we can find such a splice site, then we adjust the read alignment to match it. While this technique greatly reduces the false alignments around the splice sites, it does not eliminate the presence of spurious false exons introduced by random sequencing insertions. Pruning edges that are not supported by a minimum number of spliced reads, as described in the previous Methods section, eliminates some of the false positive edges. However, in regions of very high within-transcript sequence coverage, there may still be too many spurious nodes and edges in the splicing graph, which in turn may can cause StringTie1 to hang indefinitely. To improve StringTie2’s efficiency in such cases, we designed and implemented a pruning algorithm that reduces the size of the splicing graph to a more realistic size (see Algorithm S1 in Supplemental Material). This algorithm removes edges in the graph starting from the edge least supported by reads to the most supported edge, until the number of nodes in the splicing graph falls under a given threshold (by default 1000 nodes). Pruning edges in the splicing graph will also change the internal representation of the long reads affected by the pruning. For instance a long read that spans a node that is no longer part of the splicing graph might be represented as an interrupted read instead of a one continuous read, similar to how two paired reads are represented (see Supplementary Figure S3.b).

### Super-Read Construction and Quantification

Super-reads were constructed using code adapted from the MaSuRCA assembler (Zimin, Marcais et al. 2013). MaSuRCA builds a k-mer lookup table out of every k-mer in the input reads. It uses this to create “k-unitigs”, which are defined as sequences of maximal lengths such that every k-mer except the first and last have a unique preceding and following k-mer. Super-reads are then constructed by matching each k-mer at the ends of each short read to a unique k-unitig, effectively extending the short read as far as there is a unique extension. Note that it is possible for a super-read to contain multiple short reads, and for a short read to be contained in multiple super-reads. Not all short reads are assigned to a super-read, so both super-reads and unassigned short reads are used for assembly.

Prior to the construction of the k-mer lookup table, MaSuRCA uses QuorUM (Marcais, Yorke et al. 2015) to correct errors in the short reads. The built-in parameters that it uses for genome assembly are not optimal for transcriptome assembly. For example, these parameters include a minimum number of times a k-mer must appear to be considered high-quality, which is appropriate for genome assembly where all sequences should be covered uniformly, but not for transcriptome assembly, where some transcripts may have coverage as low as a single read. Therefore we modified these routines to remove the minimum k-mer count thresholds used for error correction. There are also certain cases where the first and/or last *k-1* bases of a super-read can extend into alternatively spliced exons, which could mislead the assembly process. To alleviate this problem, StringTie2 ignores the first and last *k-1* bases of aligned super-reads.

Because many reads may be collapsed into a single super-read, StringTie2 needs a coverage estimate with every super-read in order to calculate the expression level of any transcript with super-reads aligned to it. To estimate coverage, we first find every super-read containing each short read by matching the k-unitigs. A read is assigned to a super-read if its k-unitigs are contained in the super-read in the same continuous order (or reverse order for the opposite strand), which happens if and only if the read (or its reverse complement) is an exact substring of the super-read. During this step, we only consider super-reads that have been aligned to the reference genome. After read assignment, we use an expectation-maximization algorithm to estimate coverage for each super-read. The initial estimate sums the coverage of each read or fragment uniquely assigned to one super-read. Each iteration then recomputes coverage for every super-read by distributing coverage from each read proportionally to the previous super-read coverage estimate. This is analogous to how StringTie2 distributes coverage between transcripts. We report the computed coverage for each super-read using a special tag in the SAM output file, which is then merged with an aligned short-read SAM file which can be input to StringTie2, which uses the super-reads to weight the paths that they match in the splice graph.

## Availability

StringTie2 is implemented in C++ and is freely available as open source software at https://github.com/mpertea/stringtie2.

## Acknowledgements

This work was supported in part by grants DBI-1458178 and DBI-1759518 from the National Science Foundation, and grant R01-HG006677 from the U.S. National Institutes of Health.

