## Supplemental Material for "Transcriptome assembly from long-read RNA-seq alignments with StringTie2"

Supplementary Material

**Algorithm S1.** ***Splicing graph pruning algorithm***

**Input**.

- splicing graph $G=(V,E)$ where $V$ and $E$ represent the set of nodes and edges in the graph, respectively, and $n_i$ comes before $n_{i+1}$ on the genomic sequence *for* $\forall i \leq |V|-1$, $V=\{n_1,n_2,\ldots,n_{|V|}\}$
- an upper threshold $m$ on the number of nodes
- $W=\{w_e\}_{e\in E}$ a set of coverages of all edges in $E$.

**Output**. Pruned graph $G^p=(V^p,E^p)$ with $|V^p| \leq m$

**Algorithm**.

1. Sort edges in $G$ such that if $E=\{e_1,e_2,\ldots,e_{|E|}\}$ then $w_{e_{i+1}} \geq w_{e_i}$ $\forall i \leq |E|-1$
2. $i=0$, $V^p=V$, $E^p=E$
3. *while* $|V^p| > \mathrm{m}$ *do*
   a. remove edge $e_i$ from $E^p$
   b. *for* $n=\{n_s, n_e\}$ where edge $e_i$ links nodes $n_s$ and $n_e$ do

      *if* $n$ has no edges connecting it to other nodes in $G^p$

      remove node $n$ from $V$; $|V^p|$--

      *else if* $n_s$ and $n_{s+1}$ are adjacent on the genomic sequence (there is no intron to separate them) and there are no edges linking $n_s$ and $n_j$, $s+1<j$, *then*

      merge nodes $n_s$ and $n_{s+1}$; $|V^p|$--

      *if* $n_{e-1}$ and $n_e$ are adjacent on the genomic sequence (there is no intron to separate them) and there are no edges linking $n_j$ and $n_e$, $j<e-1$, *then*

      merge nodes $n_{e-1}$ and $n_e$; $|V^p|$--

   *i++*

   *end while*

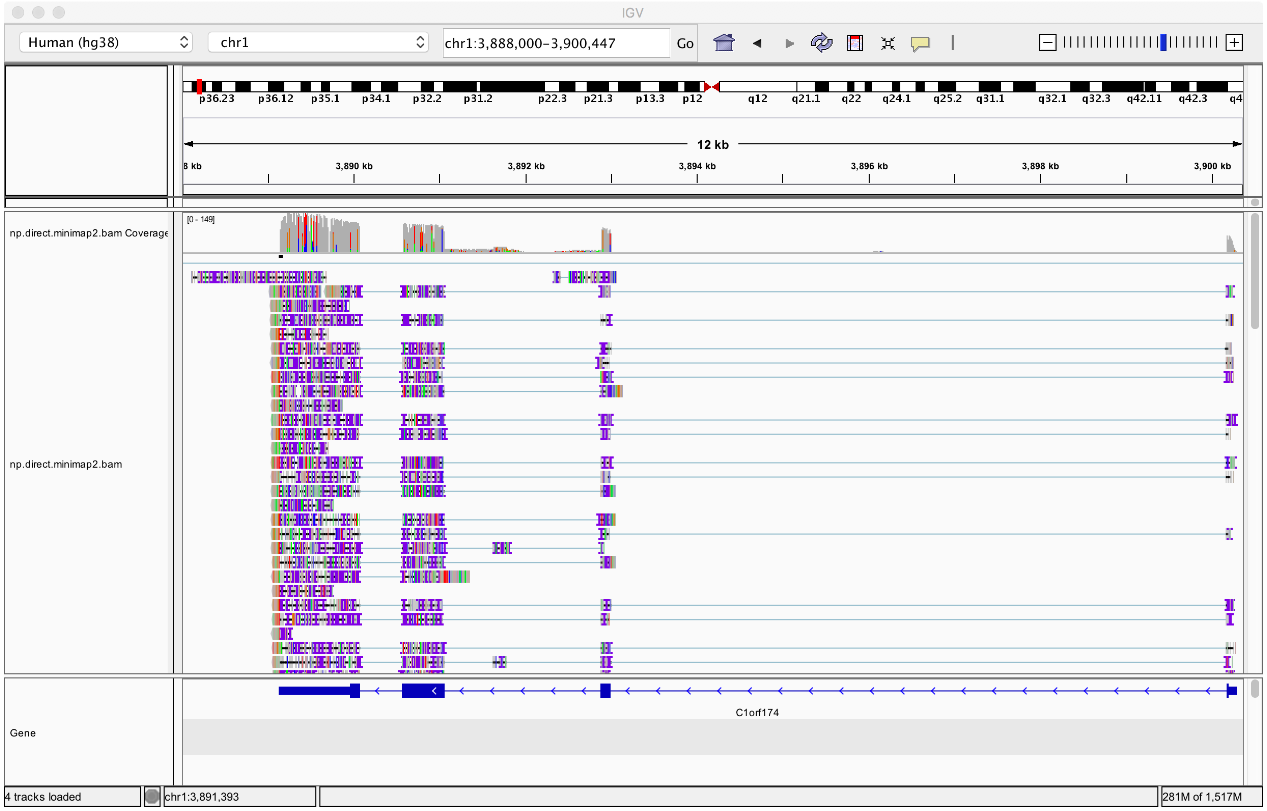


**Supplementary Figure S1.** An IGV snapshot of a aligned ONT direct RNA reads in the region of gene C1orf174. The alignments show many disagreements around the splice sites from C1orf174.

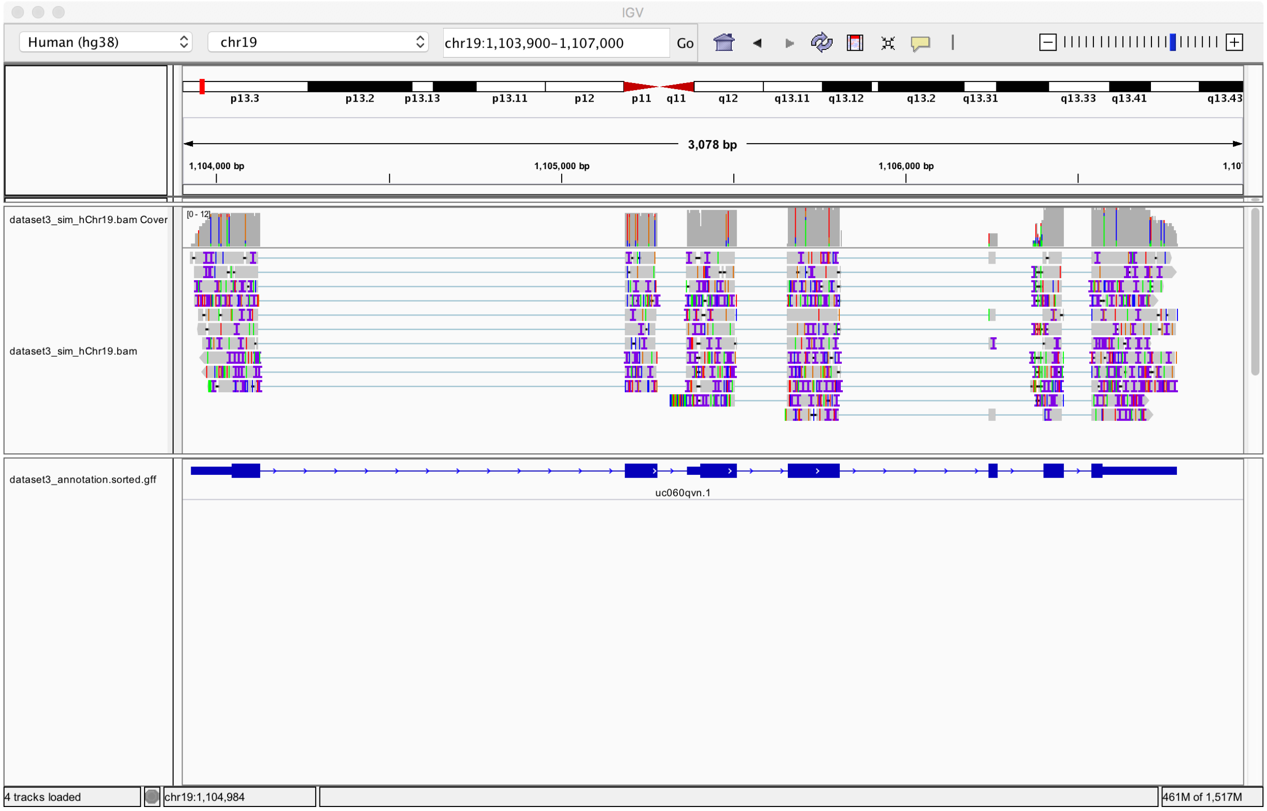


**Supplementary Figure S2.** An IGV snapshot of aligned simulated PacBio RNA reads. All reads are simulated from transcript uc060qvn.1, pictured. Some of the alignments skip the 5$^{th}$ exon in the transcript, and most alignments disagree around the splice sites.

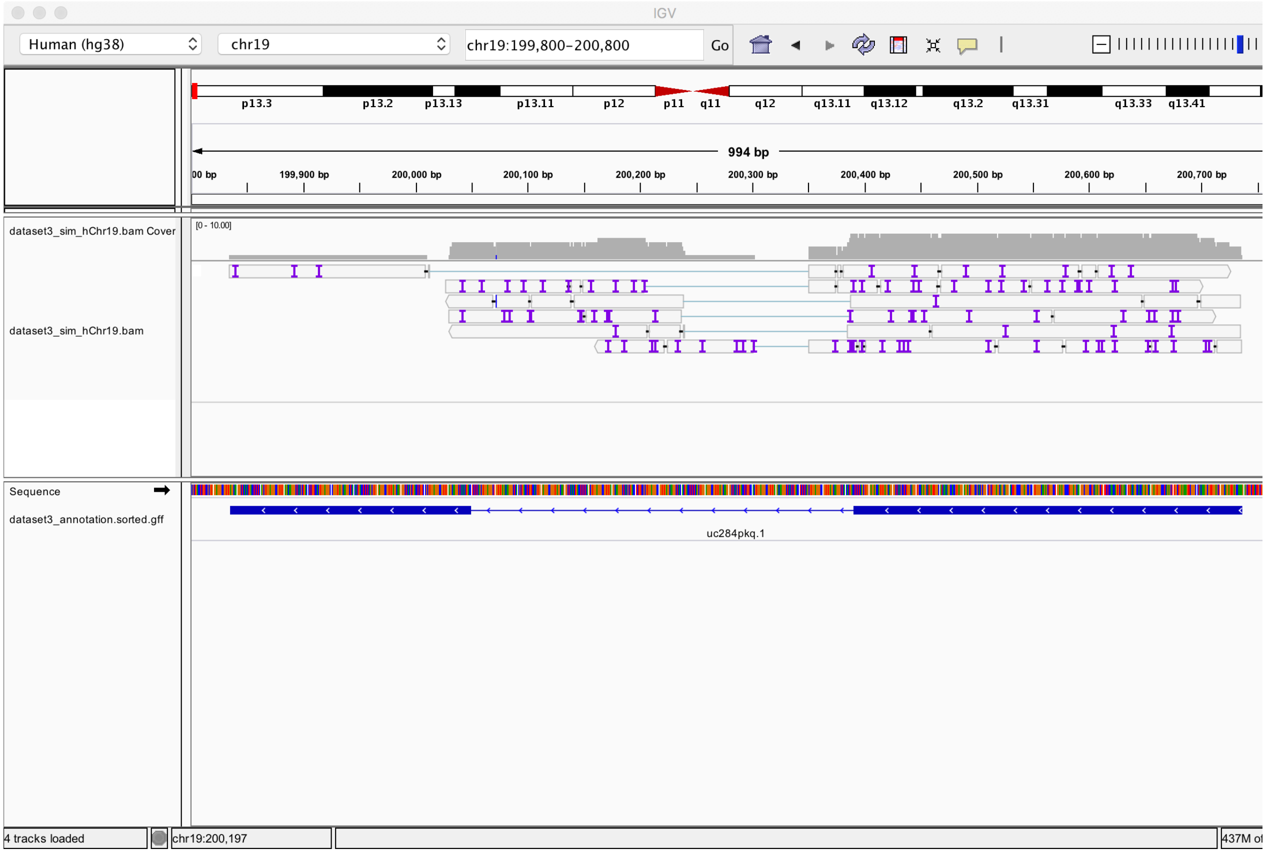


**Supplementary Figure S3.** An IGV snapshot of aligned simulated PacBio RNA reads. All reads are simulated from transcript uc284pkq.1, pictured above, and yet none of the alignments correspond to the exact exon-intron structure of the transcript.

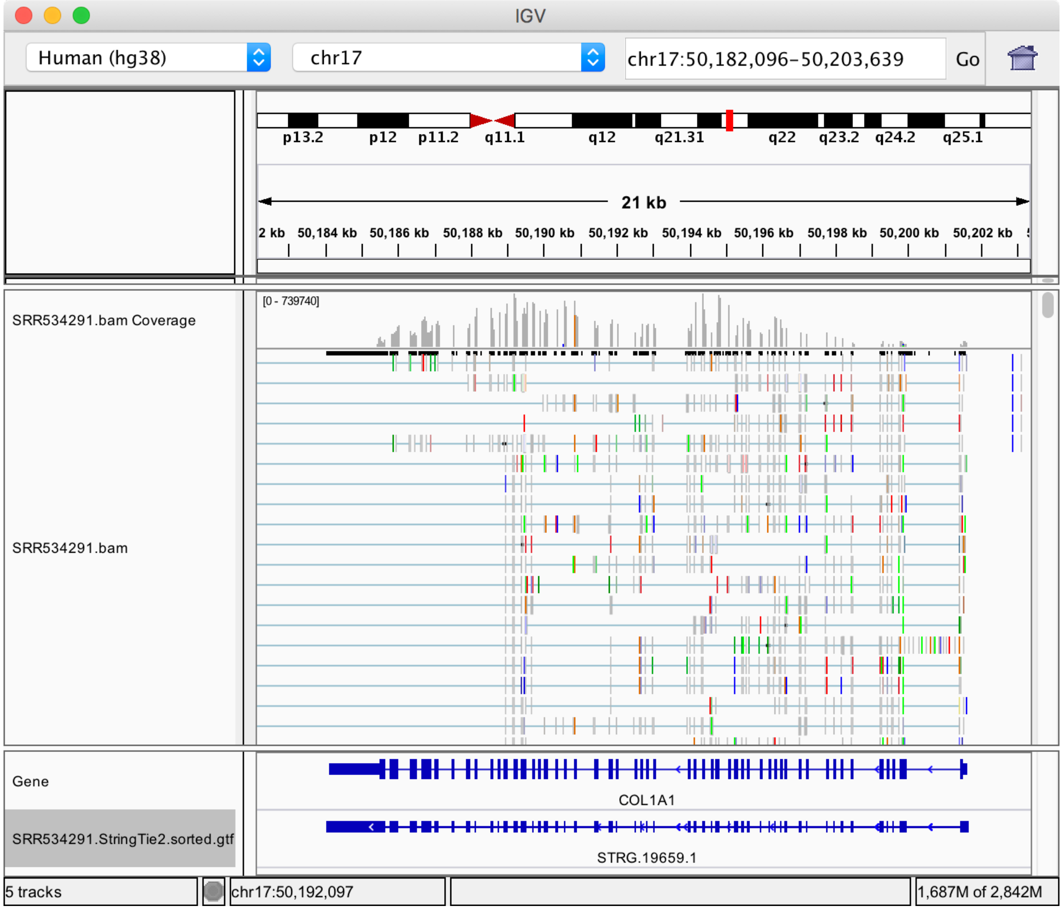


**Supplementary Figure S4.** An IGV snapshot of a highly covered transcript in the COL1A1 gene expressed captured by RNA-seq sample SRR534291 from the cytosol of fetal lung fibroblasts (GEO accession GSM981244).

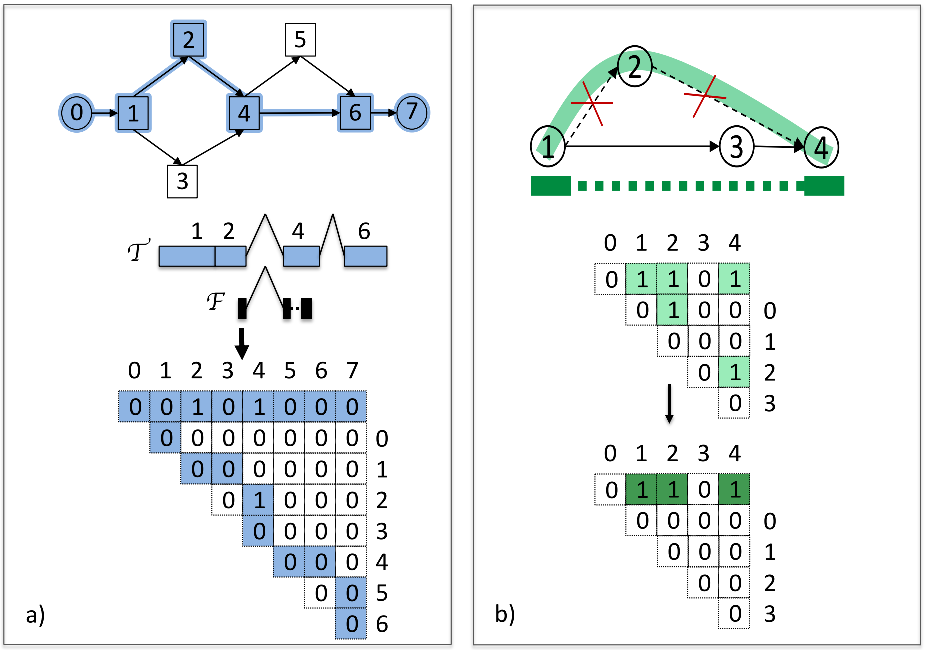


**Supplementary Figure S5. a)** Representations of a splice graph (pictured on top), an assembled transcript $\mathcal{T}'$ from the splice graph (pictured in blue as a path in the splice graph and as an exon-intron structure below the splice graph), a fragment $\mathcal{F}$ with two paired reads sequenced (pictured in black), and a bit vector corresponding to the splice graph with cells highlighted in blue representing the path of the transcript $\mathcal{T}'$, and cells set to 1 representing the fragment $\mathcal{F}$. Note that the first row in the bit vector corresponds to all the nodes in the splice graph, while the cells *(i,j)* in the matrix bellow correspond to edges that link nodes *i* and *j* in the splice graph. B). **b)** Example of a long read before (in light green), and after (in dark green) pruning the splice graph of edges (1,2) and (2,4). The read spans three nodes in the graph before pruning (1, 2, and 4), while after pruning the two edges the long read appears as a paired read instead of a continuous read and spans only two nodes (1 and 4). The bit vector representations below show the bits set before, and after pruning.

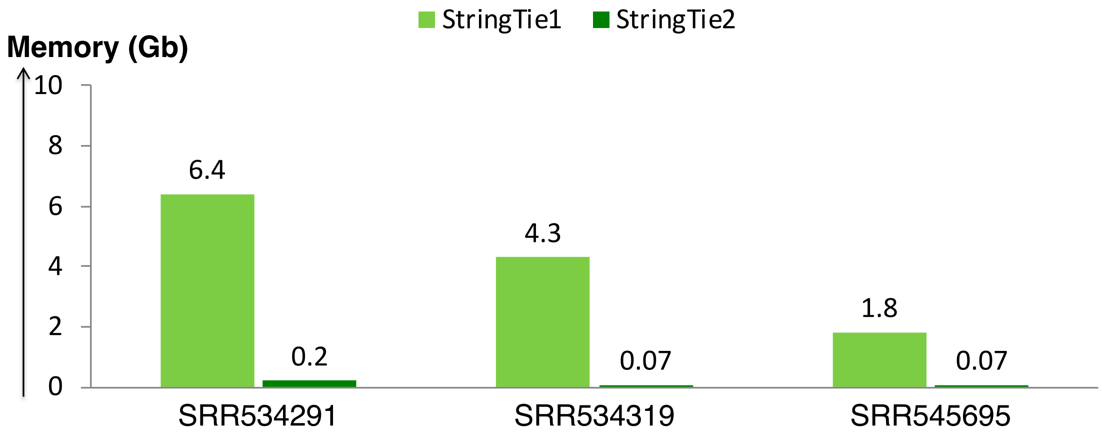


**Supplementary Figure S7.** Memory usage for StringTie1 and StringTie2 on the three datasets also included in the original release of StringTie.
